# SynTracker: a synteny based tool for tracking microbial strains

**DOI:** 10.1101/2021.10.06.463341

**Authors:** Hagay Enav, Ruth E. Ley

**Affiliations:** Department of Microbiome Science, Max Planck Institute for Developmental Biology, Tübingen, Germany

## Abstract

In the human gut microbiome, specific strains emerge due to within-host evolution and can occasionally be transferred to or from other hosts. Phenotypic variance among such strains can have implications for strain transmission and interaction with the host. Surveilling strains of the same species, within and between individuals, can further our knowledge about the way in which microbial diversity is generated and maintained in host populations. Existing methods to estimate the biological relatedness of similar strains usually rely on either detection of single nucleotide polymorphisms (SNP), which may include sequencing errors, or on the analysis of pangenomes, which can be limited by the requirement for extensive gene databases. To complement existing methods, we developed SynTracker. This strain-comparison tool is based on synteny comparisons between strains, or the comparison of the arrangement of sequence blocks in two homologous genomic regions in pairs of metagenomic assemblies or genomes. Our method is executed in a species-specific manner, has a low sensitivity to SNPs, does not require a pre-existing database, and can correctly resolve strains using complete or draft genomes and metagenomic samples using <5% of the genome length. When applied to metagenomic datasets, we detected person-specific strains with an average sensitivity of 97% and specificity of 99%, and strain-sharing events in mother-infant pairs. SynTracker can be used to study the population structure of specific microbial species between and within environments, to identify evolutionary trajectories in longitudinal datasets, and to further understanding of strain sharing networks.

## Introduction

Strains of the same microbial species (conspecific strains) often present large phenotypic differences, despite having very similar genotypes (Van Rossum et al. 2020). Previously published examples for phenotypic differences between strains of the same species include pathogenicity (Pierce and Bernstein 2016), commensalism (Leimbach, Hacker, and Dobrindt 2013), drug response (Maier et al. 2018) and susceptibility to infection by phages (Holmfeldt et al. 2007). In host associated microbial communities, species can stably coexist for years (Faith et al. 2013; Lloyd-Price et al. 2017), potentially evolving into host-specific strains (S. Zhao et al. 2019). Occasionally, host specific strains could be transferred to or from other hosts, along familial networks (Yassour et al. 2018), through the built environment (Brooks et al. 2017) or following fecal microbiota transplantation (Li et al. 2016; Smillie et al. 2018). The ability to identify and follow conspecific strains is required to understand the mechanisms of between-host strain transmission, within-host evolution, and how these forces interact to shape microbial communities.

Methods to track strains using short-read data currently belong to one of two main classes: *de-novo* assembly of contigs from metagenomes, and methods relying on alignment of genomic sequences to a reference database (reviewed in (Anyansi et al. 2020)). Methods in both classes usually rely on detection of single nucleotide polymorphisms (SNPs). In assembly-based methods, high sequencing depth is required to overcome sequencing errors and natural variation in the population, making these tools more suitable for the study of low-complexity microbial communities (Anyansi et al. 2020). Moreover, identifying SNPs based on metagenomic assembled genomes (MAGs) can introduce errors in low quality MAGs (Van Rossum et al. 2020). On the other hand, methods based on comparisons to a reference database usually require lower sequencing depth, although SNP detection in these methods can be limited by natural variation in the population and the degree of similarity between the community members and the reference genome (Bush et al. 2020). Moreover, reference-based methods can only track strains belonging to well-studied species, for which a suitable reference database has been generated.

To complement methods relying on SNP information, we developed SynTracker, an approach to identify and track closely-related strains using genome microsynteny (the local conservation of genetic-marker order in genomic regions). Gene synteny (organization of genes along two chromosomes) has been used to estimate evolutionary distances between genomes (Lemoine, Lespinet, and Labedan 2007; Alexeev and Alekseyev 2017; T. Zhao et al. 2021) and to identify horizontal gene transfer events (Adato et al. 2015). SynTracker uses pairwise comparisons of homologous genomic regions in either metagenomic, or genomic assemblies, followed by scoring the average synteny per pair of strains. SynTracker is relatively insensitive to SNPs, and requires only a single reference genome per species (either complete or draft and with no regard to its annotation level). Here, we apply SynTracker to compare the within-population synteny of *in-silico* evolved bacterial populations and to reproduce known within-species phylogenies of *E. coli*, using a fraction of the entire genome length. Additionally, we define a synteny-score cutoff to identify strains residing in the same hosts’ gut microbiomes over time. Finally, we apply SynTracker to a gut microbiome metagenomic dataset consisting of samples obtained from mothers and their infants (Bäckhed et al. 2015), and describe a high degree of strain sharing between mothers and their infants in species which colonize the infant gut.

## Results

### Pipeline description

SynTracker is based on the identification of synteny blocks in pairs of homologous genomic regions derived from isolate genomes, metagenomic assemblies or metagenome-assembled genomes (MAGs). The pipeline accepts as input a reference genome per species of interest, either fully or partially assembled, and a collection of metagenomic assemblies (or genomes, if single genomes are to be compared). In the first step of the pipeline (Figure 1A, ***methods***), the reference genome is fragmented to create a collection of 1 kbp genomic regions, located 4kbp apart (“central-regions”). Next, we convert the collection of per-sample metagenomic assemblies (or genomes) to a BLAST (Altschul et al. 1990) database and use the central-regions as queries for a high stringency *BLASTn* search (Identity=97%, minimal query coverage=70%) to minimize the possibility of receiving either multi-species hits or hits located within regions with high copy number variation. Next, For each BLAST hit we retrieve the target sequence and the flanking 2 Kbp regions upstream and downstream of the target sequence. This strategy results in high specificity when identifying homologs to the central regions, while allowing for high variance in the sequence composition of the flanking regions.

**Figure 1:**
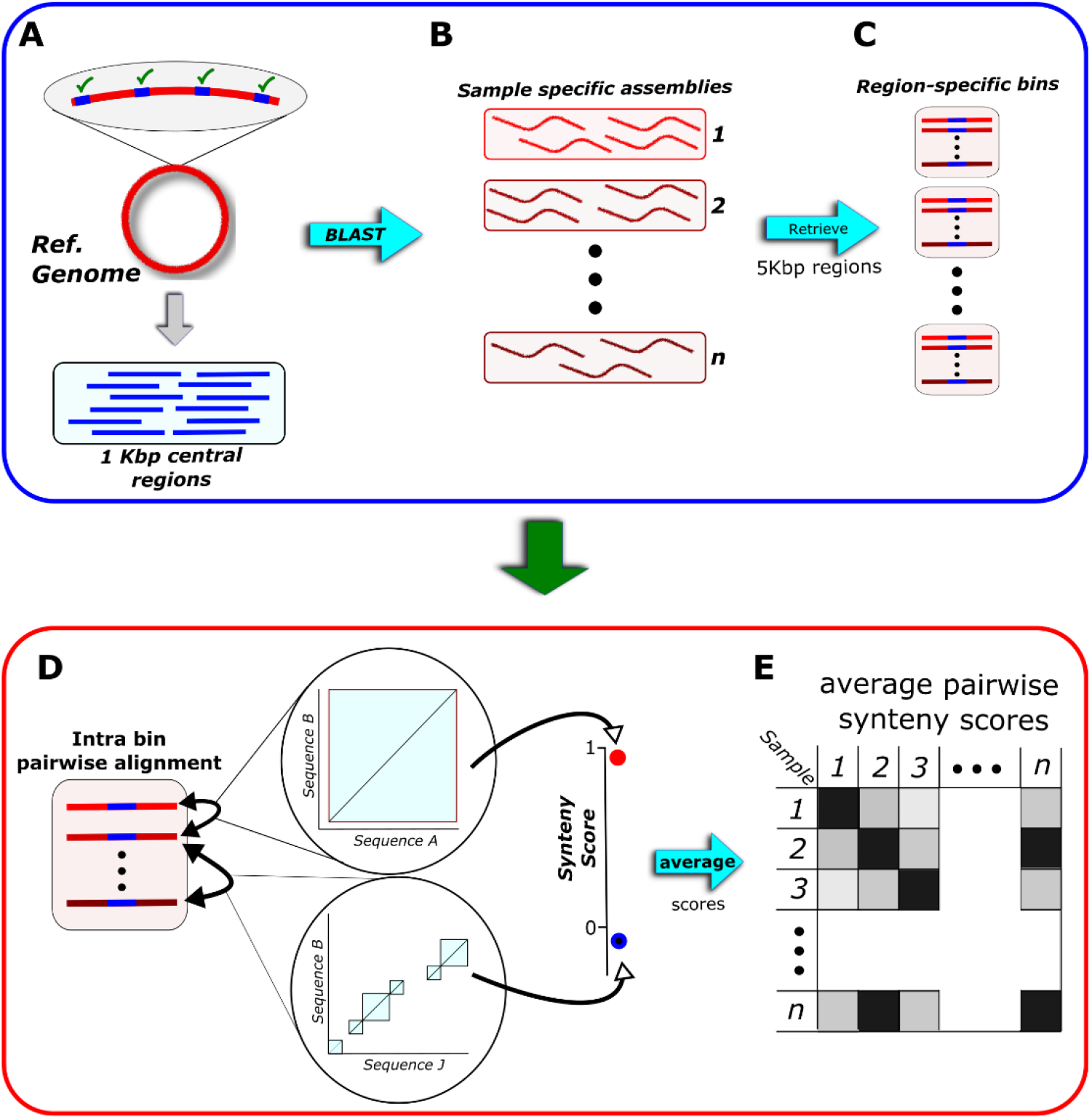
Illustration of the SynTracker pipeline. **A.** The reference genome is fragmented to yield “central-regions”, i.e., 1 kbp long regions located 4kbp apart. **B**. Each central region is used as a query for a BLAST search against a collection of sample-specific assemblies (or genomes, if isolates are analyzed). **C.** BLAST hits are retrieved with 2 kbp on each side of the hit. All bins resulting from the same BLAST search are placed in the same Region-specific bin. **D**. Within each bin, all-versus-all pairwise alignment is performed to identify synteny blocks in pairs of sequences. Synteny scores are calculated based on the number of blocks and their accumulative length. **E**. For each pair of samples (or genomes) *n* regions are sampled and their synteny scores are averaged to yield the average pairwise synteny score (APSS).

Next, each collection of homologous ~5Kbp regions (*i.e*., derived from a BLAST search using the same central-region query) is assigned to a region-specific bin (Figure 1B). Within each bin we perform an all vs. all pairwise sequence alignment to identify synteny blocks using the DECIPHER R-package (Wright 2016). Then, for each pairwise alignment we calculate the region-specific pairwise synteny score (*see Methods*). This score is based on two parameters: the number of synteny blocks identified in each pairwise sequence alignment, and the overlap between the two sequences. The synteny score is inversely proportional to the first and directly proportional to the second.

A single synteny block in a pairwise alignment can stem from two genomic regions with a high sequence similarity. A high number of synteny blocks can result from insertions, deletions, recombination events or several SNPs located within a close proximity in just one of the two sequences. The sequence overlap is defined as the ratio of the accumulative length of all blocks to the length of the shorter DNA region in each pairwise comparison. The region-specific pairwise synteny has a maximal value of 1, reflecting identification of a single synteny block and overlap of 100% (Figure 1C). After calculating the per-region synteny scores in all bins, we randomly subsample *n* regions per a single comparison of metagenomic samples (or genomes), and determine the Average Pairwise Synteny Score (***APSS***, Figure 1D).

### Analysis of in-silico evolved bacterial population reveals low sensitivity to SNPs

While numerous tools to study conspecific strains are available, most rely on SNP data (Anyansi et al. 2020). Since the synteny approach was designed with the aim of complementing existing methods, we minimized the effect of SNPs on the APSS values. Our approach was designed to give a higher weight to insertions, deletions and recombination events, which are less abundant than SNPs, and are less likely to result from sequencing errors (Schirmer et al. 2016).

To examine the performance of our approach and estimate the effect of different genomic variations on the synteny scores, we used *in-silico* simulations of the evolution of bacterial populations. To generate simulated population data, we used Bacmeta (Sipola, Marttinen, and Corander 2018), a simulator for genomic evolution in bacterial metapopulations. We performed two types of simulations: in the first, the population evolved by introducing SNPs exclusively, with a frequency of 1*10^−6^ substitutions per nucleotide per generation. In the second simulation, we introduced both insertions and deletions, each with a frequency of 5*10^−8^ mutations per nucleotide per generation. In both simulations we set the population size to 10000 bacterial cells and analyzed three genomic regions, each with a length of 20 Kbp. We carried out the simulation for 3,000 generations and randomly subsampled 20 cells every 300 generations.

At each timepoint, for each genomic region, we calculated all pairwise synteny scores in addition to all pairwise sequence identities (Figure 2). In simulations using SNPs, the minimal average blast identities were 99.48%, 99.46% and 99.5%, for regions 1, 2 and 3, correspondingly. The lowest average BLAST identities in simulations based on insertions and deletions were higher, at 99.79%, 99.78% and 99.84%. In accordance with the expectation that the synteny approach is more robust to changes in SNPs than to indels, the synteny scores in SNP-based simulations were higher (0.905, 0.849 and 0.863) than in indel-based simulations (0.067, −0.0589, 0.0093). It is important to emphasize that this difference was achieved even though the mutation frequency in SNP based simulation was ×10 higher than the Indel-based simulations. The lower synteny scores of genomic regions in the indel-based simulations further support the higher sensitivity of the synteny-based approach to indels, which are used as a “genomic fingerprint” in our method.

**Figure 2:**
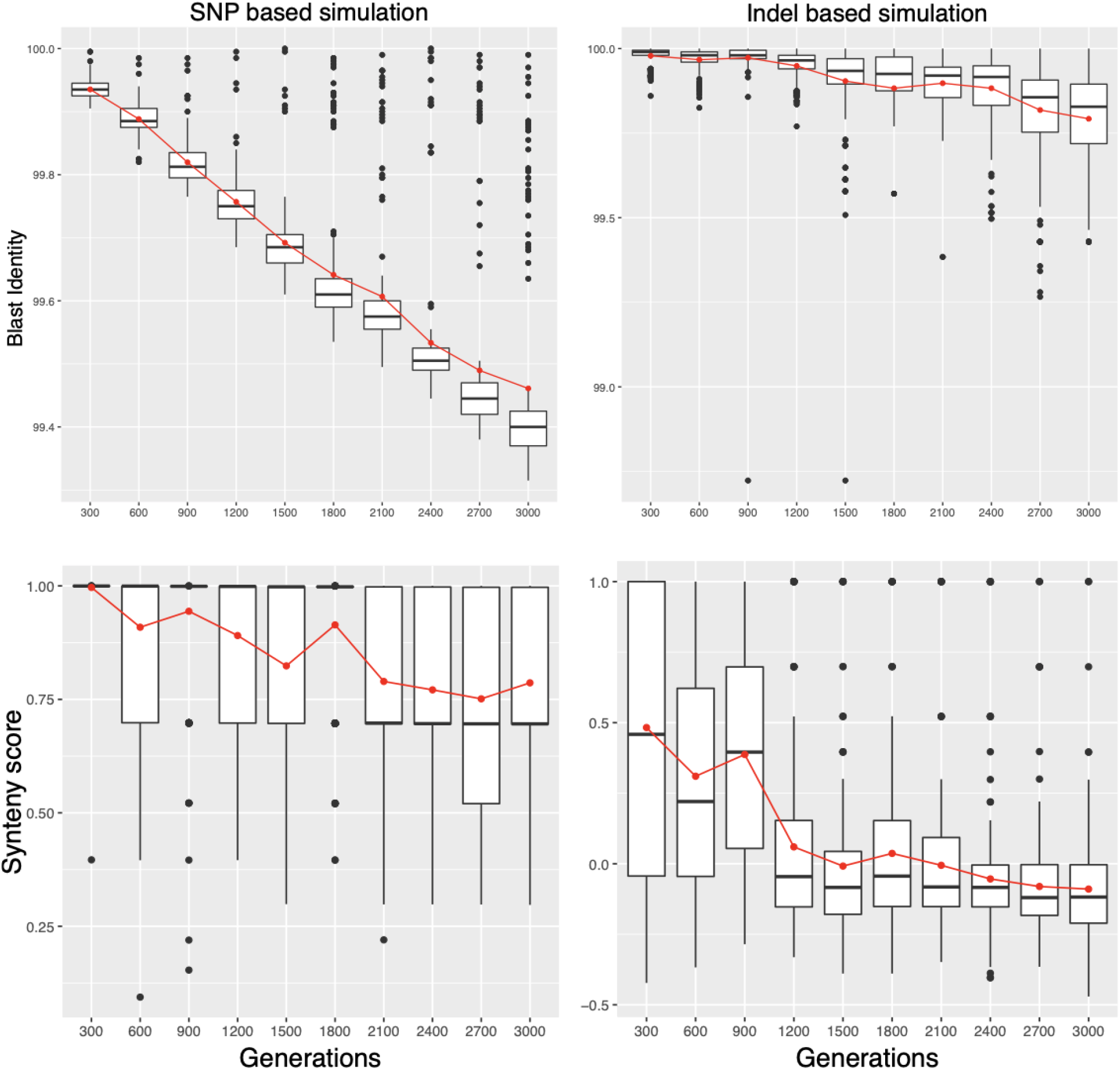
Analysis of the genomic diversity of *in-silico* evolved bacterial populations. Simulations were carried out for 3000 generations. In each time point, 20 cells were sampled, and a pairwise comparison of the same 20 kbp region was performed using BLASTn (top panels) and SynTracker (bottom panels). Horizontal black lines mark the group median, the red lines correspond to the group mean. Left plots show analyses of a representative simulation carried through the exclusive introduction of SNPs at a frequency of 1*10^−6^ substitutions per nucleotide per generation. Right panels show a simulation based on indels at a frequency of 1*10^−7^ substitutions per nucleotide per generation.

### The synteny method can reconstruct phylogenies using a fraction of the reference genome

We examined the performance of SynTracker for the comparison of closely related genomes and to use the resultant APSS values as a basis for generating phylogentic trees. We used a recently published whole genome MASH based classification of >10K *E.coli* genomes that identified 14 distinct phylogroups (Abram et al. 2021). We randomly selected 10 genomes per phylogroup, for a total of 140 *E.coli* genomes. We analyzed the set of genomes eight times, and in each iteration, we randomly selected a different number of 5 kbp regions per pairwise comparison (15-200 regions/pairwise comparison, representing ~1.4-18.5% of the *E.coli* O157:H7 genome length) to create the final matrices holding the APSS values (Figure 1D). These matrices were used to generate UPGMA phylogenetic trees based on the APSS distances (see methods). With subsampling of 200 regions/pairwise comparison, we recapitulated the classifications of 139/140 *E. coli* genomes to the published phylogenetic groups. When reducing the number of regions used per pairwise comparison to 40 (roughly equal to 3.6% of the full genome’s length) the phylogeny we obtained matched the published one, with the same taxa forming the previously designated groups, except for 4 genomes (Figure 3). These results underscore the utility of the synteny approach in the analysis and comparison of bacterial genomes, even at a very low levels of genome completeness.

**Figure 3:**
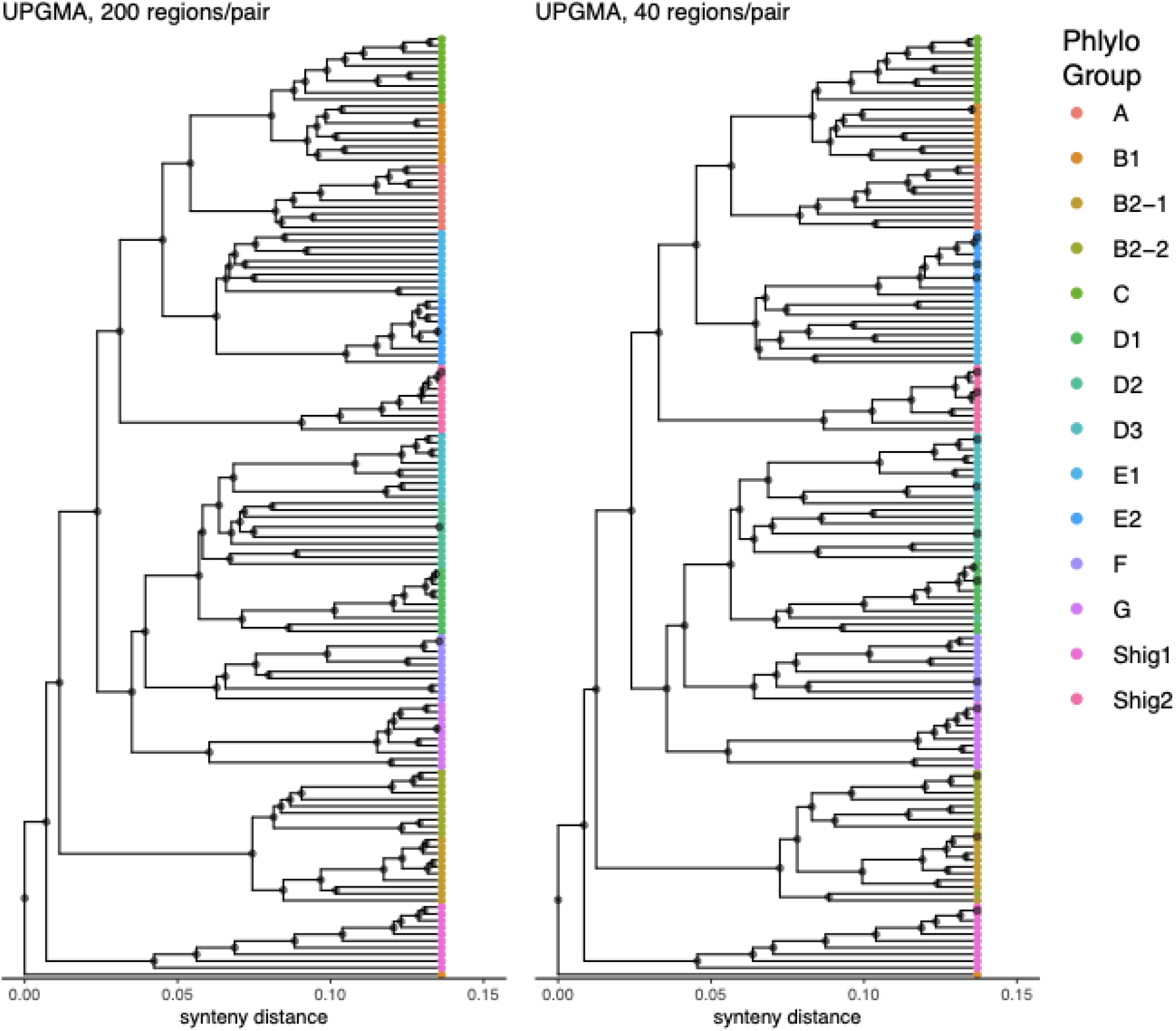
Phylogenetic trees of 140 *E. coli* genomes, belonging to 14 different phylogroups, based on Average Pairwise Synteny Scores (APSS). **A.** A tree based on 200 regions/pair (accumulative length of ~1 mbp) correctly classified 139 genomes. **B.** A tree based on 40 regions/pair shows correct classification of 136 genomes.

### Assessment of the method’s performance in identifying within-host strains

We tested SynTracker for the detection of closely-related bacterial strains within whole-community metagenomic samples. As bacterial strains can reside in the human gut for years (Schloissnig et al. 2013; Faith et al. 2013), we applied SynTracker to differentiate between within-individual bacterial strains and conspecific strains inhabiting different hosts. We calculated the APSS values of closely related strains, classified to one of 38 different bacterial species (*table s1*), in 223 gut metagenomes collected from 84 healthy westernized human donors (Poyet et al. 2019) (*table s2*).

For each of the studied species, we used a publicly available reference genome (*table s1*), which was fragmented into a collection of 1 kbp “central regions”, as described above. Next, we performed a per-sample, *de novo* metagenomic assembly to construct our “search-space” (methods, Figure 1). The metagenomic assemblies were divided randomly into training and testing sets (117 and 106 samples, obtained from 45 and 43 donors, respectively). For both sets, we calculated eight different final APSS matrices per species, after randomly selecting *n* regions per pairwise comparison (*n*=15-200 5 kbp regions; see methods). Following the calculation of the APSS matrices for each species, we classified pairwise comparisons in the training set to those originating from the same host at different time points (within-host) and those that originate from different hosts (between-host, Figure 4a). With the classification of pairwise comparisons in the training set used as ground-truth, we created a receiver operating characteristic curve (ROC) (Fawcett 2006) for each combination of species and subsampling value (Figure 4B).

**Figure 4:**
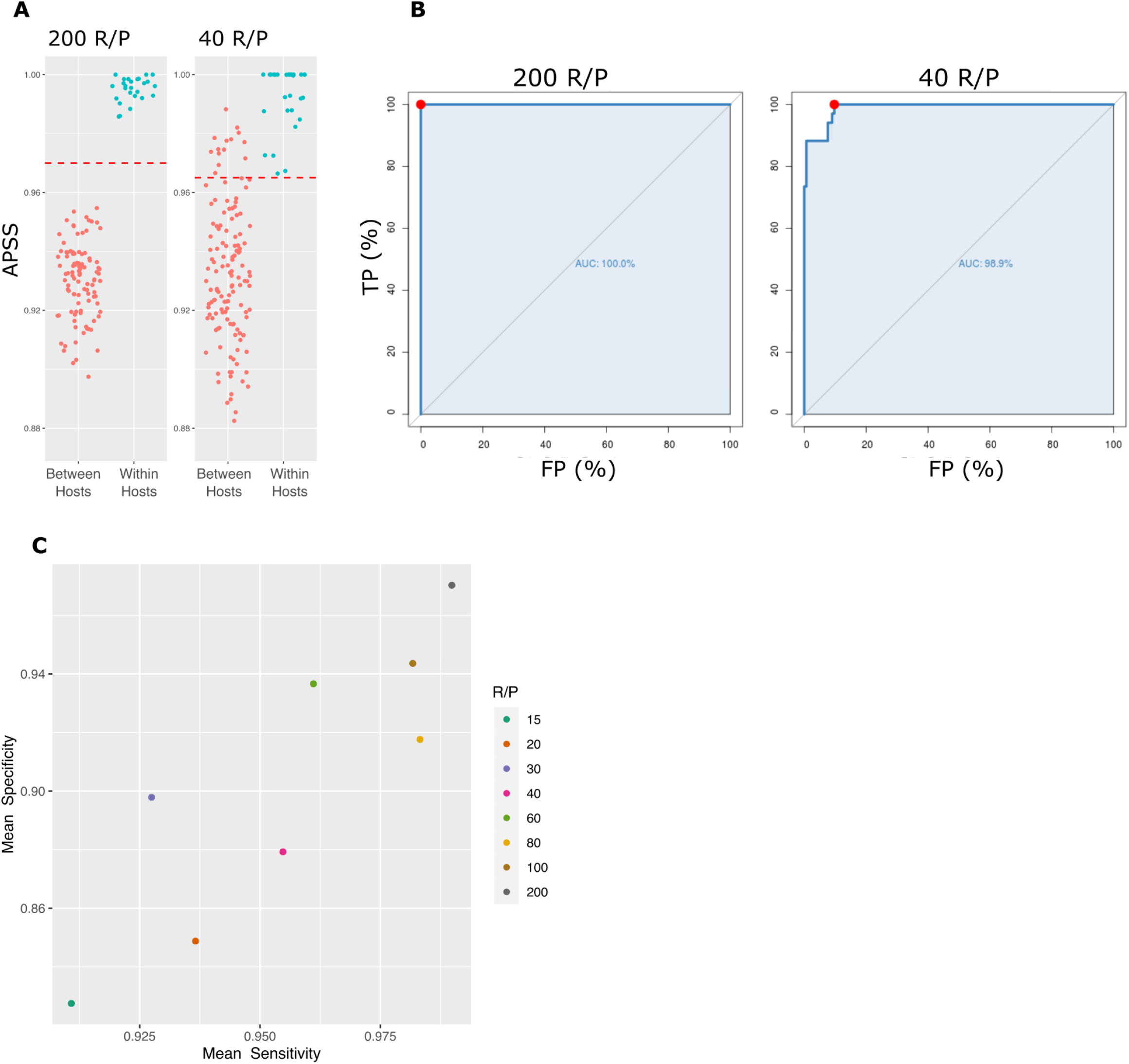
identifying within-host strains. **A**. Representative synteny-based analyses of *B. fragilis* strains in longitudinal metagenomic samples, obtained from the same host (cyan) and different hosts (pink), using 200 and 40 regions/pairwise-comparison. Dashed red lines represent the APSS threshold differentiating between strains residing in the same host and strains residing in different hosts, as calculated in B. **B**. ROC plots of the same analyses shown in A. Red dots correspond to the Youden point and give the APSS score that yields the optimal combination of specificity and sensitivity. **C.** The mean specificity and sensitivity of the synteny approach, calculated at different numbers of regions/pairwise-comparison. R/P-regions/pairwise comparison, TP-True Positive, FP-False Positive.

ROC plots are created by assessing the sensitivity and specificity (proportional to the percent of true positive and false positive observations) of a classifier while using different discrimination values, which in this analysis was APSS. To determine the APSS values that optimally discriminate between strains residing in the same host and strains identified in different hosts, we calculated the J-index (Youden 1950) for each combination of species and subsampling depth. Finally, we used these APSS thresholds to determine the specificity and sensitivity of our method, by introducing them to the testing set. Not surprisingly, we found a direct correspondence between the number of subsampled regions per pairwise comparison and the sensitivity and specificity of our method (Figure 4c, Table 1), with maximal sensitivity and specificity of 99% and 97%, for comparisons calculated using 200 regions/pairwise-comparison. While using a small number of regions/pairwise-comparison mostly results in lower accuracy, the decision to use such values may be justified by the inclusion of additional samples in the analysis. Therefore, it is up to the researcher to decide whether to prioritize increased accuracy or sample size for any given analysis.

**Table 1:**
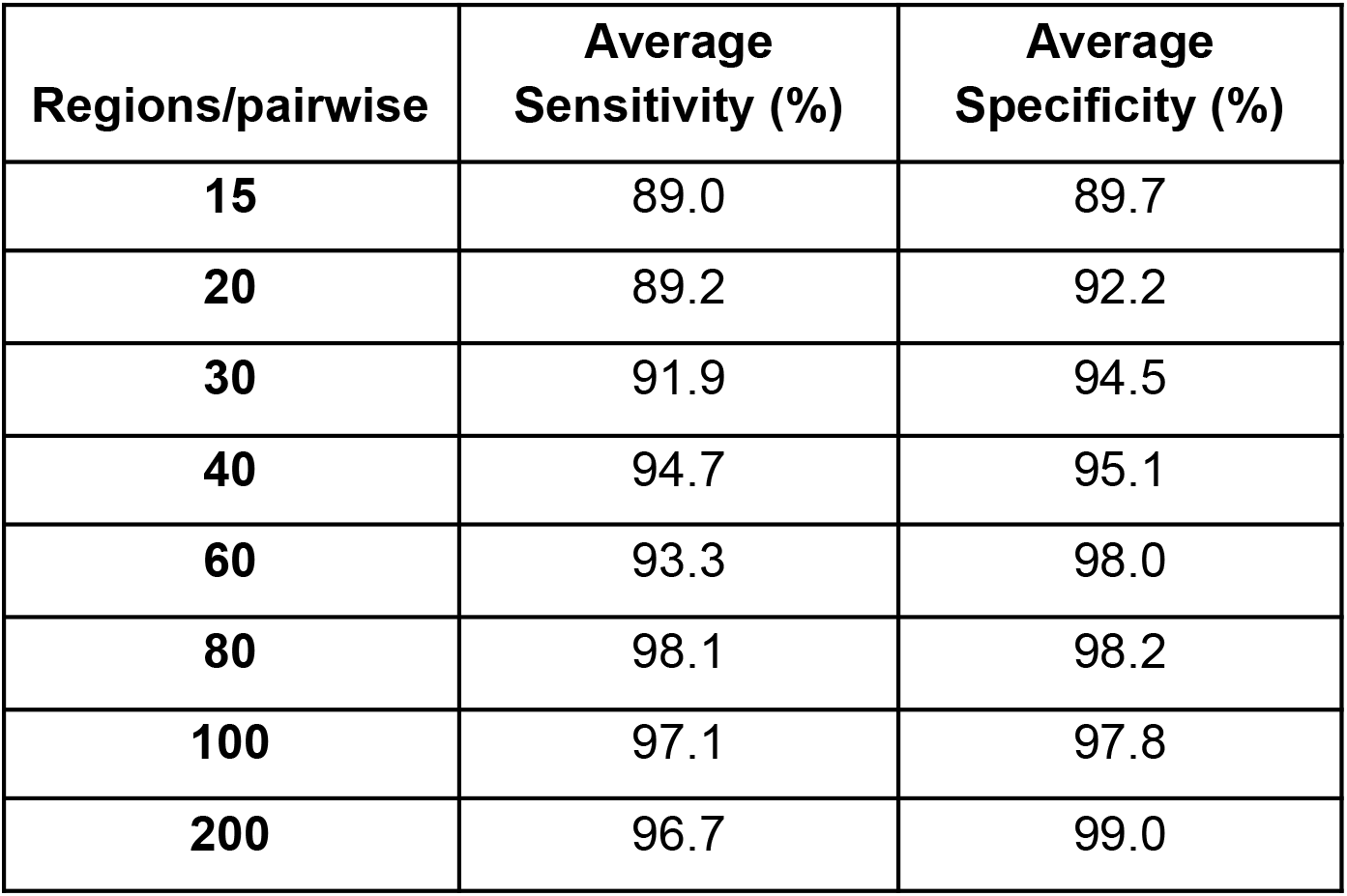
sensitivity and specificity of SynTracker using different numbers of genomic regions per pairwise comparison.

### Identifying mother-infant strain transmission

After verifying that the synteny method can be used to track host-specific strains in human gut metagenomes with high accuracy, we used this method to identify strain-sharing events between hosts. Our overarching goal was to study the role of vertical strain transmission (i.e., transmission from mother to infant) in the colonization of the human gut at early infanthood. Previously, vertical strain transmission was studied using culture-based techniques (Milani et al. 2015; Makino et al. 2013), which are inherently limited to specific taxa. More recently, vertical strain transmission was studied by identifying SNP profiles in metagenomic data(Nayfach et al. 2016; Yassour et al. 2018).

To study mother-infant strain transmission using strain synteny, we analyzed the dataset collected by Bäckhed and colleagues (Bäckhed et al. 2015). These data contain stool-derived metagenomes obtained from 98 mothers and their infants, sampled at ages of 4 days, 4 months and 1 year post-birth. We assembled the metagenomic samples de-novo (see *methods*) and calculated the APSS scores, for a collection of 38 bacterial species (Figure 1). In order to maximize the number of samples included in per-species analyses, we used 30 regions per pairwise comparison.

We expected the gut microbiome of 4-day old infants to contain strains vertically transmitted from their mothers. Therefore, we predicted that true mother-infant-pairs (MIP) will show significantly higher APSS values compared to unrelated MIP, in comparisons of mothers and newborn infants. Therefore, for each analyzed species, we grouped mother-infant comparisons by the relatedness of the pair and compared the APSS values of the two groups (*i.e*., true MIP and unrelated MIP). We only considered species with at least 6 pairwise comparisons as suitable for statistical hypothesis testing (Wilcoxon rank test, single tailed, Benjamini-Hochberg multiple testing corrected (Benjamini and Hochberg 1995)). Only a small subset of 9 species passed our criterion in the newborn age group, which could be explained by the low complexity of the newborn gut microbiome and is in agreement with previous findings regarding the maturation of the infant gut microbiome (Koenig et al. 2011). As expected, most species in this age group (7/9) had significantly higher APSS values in true MIP (q-value<0.05, figure 5). Additionally, 80/84 of the strain comparisons in true MIP had APSS value > 0.94, and therefore were considered as a vertical strain transmission, based on the APSS threshold described above.

**Figure 5:**
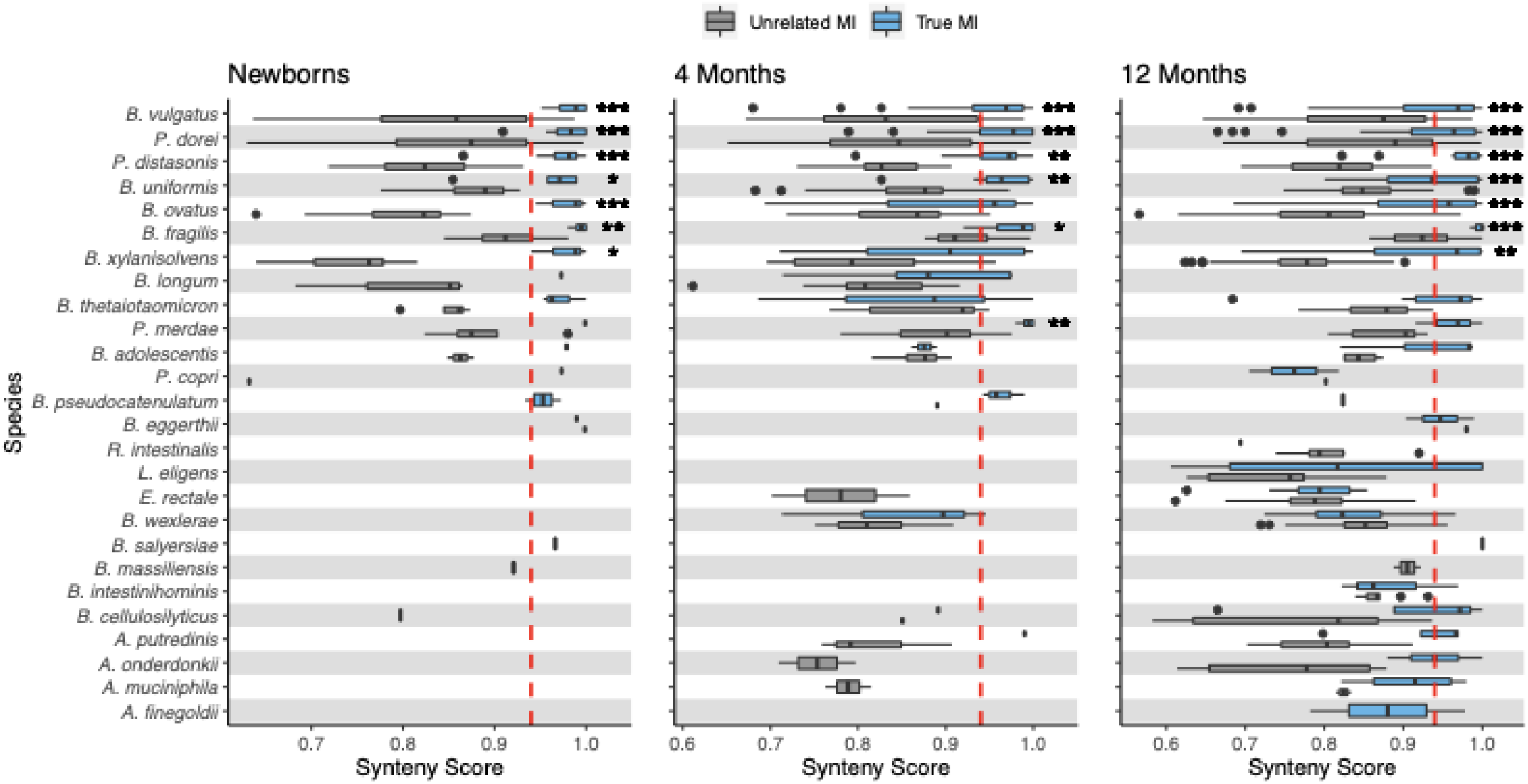
Vertical strain transmission in Mother-Infant Pairs. The Synteny of strains residing in mothers and their infants at ages 4 days, 4 months and 1 year (True MI, blue boxes) was compared with the synteny of strains in unrelated mothers and infants (unrelated MI, grey boxes). Comparisons to the right of the dashed red lines are considered as vertical transmission events. Stars correspond to Benjamini-Hochberg corrected p-values (Wilcoxon-Mann-Whitney test): q-value * <5×10^−2^, ** < 5×10^−3^, *** <5×10^−5^.

We observed that in the 4- and 12-month age groups, strains of species identified in the newborn group remained similar to those of the mothers, in true MIP (adjusted p-value < 5*10^−4^ in the 12-months group, Figure 5). In contrast, late colonising species, identified only in later samples, did not have significantly higher APSS values in true MIP compared to unrelated MIP. Overall, in the 12-months old group, we compared 240 strains in true MIP pairs, out of which 126 could be considered as resulting from a strain-sharing event.

## Discussion

In this report we introduce SynTracker, a method for tracking closely related microbial strains using genome synteny. SynTracker requires as input a collection of genomes or per-sample assembled metagenomic contigs, a reference genome file, and a metadata file. The SynTracker code can be used as a standalone tool or as a part of a custom pipeline.

We designed SynTracker to complement other existing methods that track closely related strains, which mostly rely on SNP profiles or on analysis of specific sets of genes. In studies tracking strains across individuals, reliance on SNP information could be potentially limiting: environmental changes can spur the emergence of hypermutator strains (Travis and Travis 2002; Swings et al. 2017), increasing the point-mutation rate by a factor of up to 150-fold (Wielgoss et al. 2013). To avoid the limitations of these tools, we intentionally set the default pairwise-alignment parameters in our pipeline to have low sensitivity to SNPs, which may be erroneously identified due to sequencing and amplification errors. When examining SynTracker’s performance using *in-silico* evolved bacterial populations, we observed that, as expected, populations that evolve exclusively through introduction of SNPs had a marginal reduction in the synteny scores compared to populations that evolved through introduction of insertions and deletions at a lower mutation frequency. This characteristic of our approach makes it a good candidate to complement existing SNP-based tools and also makes it ideal to track closely related strains in data produced using long-read sequencing methods, as the error rate of these methods is higher than in methods based on short-reads (Amarasinghe et al. 2020).

While some popular tools for conspecific strain analysis require a pre-existing gene database, our approach requires only a single reference genome per species, either fully assembled or as a collection of contigs. This feature is advantageous, as it allows for tracking strains of relatively understudied species. As state-of-the-art methods for assembly of genomes from metagenomes yield ever larger collections of MAGs, we propose a potential workflow, in which the MAG collection is clustered to create “species representative genomes”, which could be used as the reference genome in our pipeline. This approach, which is also utilized in the InStrain program (Olm et al. 2021), can expand our ability to study strains of novel species.

One of the most important assets of our approach is its ability to track closely related strains using only a small fraction of the full length of the genome. We were able to reconstruct the phylogeny of 140 *E.coli* strains using <20% of the length of the reference genome and to identify within-host strains with an average sensitivity of 97% and specificity of 99%, using the same accumulative length of the compared regions. The ability to track strains using a fraction of the full genome length is especially important when analyzing MAGs with low completeness values or less abundant taxa in metagenomic assemblies.

We examined the performance of our method by identifying strains residing in the same human hosts over time periods of a few weeks to two years. We observed a decrease in the performance of the approach with the reduction in the number of regions used per pairwise comparison. On the other hand, reducing the number of regions could increase the number of samples included in the final analysis. The SynTracker pipeline provides a number of average-synteny score tables, prepared using 20-200 regions/pairwise-comparison. It is up to the user to select the relevant table, based on their specific needs and dataset.

When investigating low abundance taxa, metagenome assembly might yield relatively short contigs. In those instances, the likelihood of identifying a sufficient number of overlapping 5 kbp regions in any given two metagenomic assemblies is reduced. In such cases, it could be beneficial for the user to perform the synteny-based analysis on shorter genomic regions. This could be easily achieved by reducing both the length of the flanking-regions and the spacing between the “central-regions” (Figure 1A, methods).

To demonstrate the use of SynTracker we analyzed the metagenomic dataset collected by Bäckhed et al (Bäckhed et al. 2015), who followed a cohort of mothers and their infants from birth to one year of age. Since SynTracker uses pairwise comparisons of homologous genomic regions, the number of pairwise comparisons increases exponentially with increasing numbers of samples. To reduce the overall running time of our pipeline without losing relevant information (*i.e*., comparisons of true MIP and comparisons of longitudinal samples), we divided the dataset into 20 bins, while keeping all same-family samples in the same bin. Using this strategy, we were able to reduce the number of pairwise comparisons by a factor of ~21 at the cost of losing some between-family comparisons, which were only used in our analysis as a control group, relative to the true MIP group. We strongly recommend this strategy to researchers analyzing larger datasets consisting of hundreds of metagenomic samples and dozens of reference genomes.

Our analysis of the Backhed et al. dataset showed that early colonizing species, that inhabit the guts of both the mothers and the newborn infants, had higher APSS values in comparisons of MIPs, compared to unrelated MIPs. Moreover, a striking majority of the strains tracked in true MIPs (newborn age group) had APSS scores high enough to be considered within-host strains. In the 12-month-old group, early colonizing species maintained the higher APSS values in true MIPs, compared to unrelated MIPs, while no significant difference was observed for late colonizing species. These results suggest that vertical strain transmission plays a role in the acquisition of early colonizing species, while late colonizers could be obtained from additional sources as well.

## Conclusions

We have introduced SynTracker, a tool for tracking conspecific strains and to evaluate their relatedness using genome synteny, in both genomes and metagenomic assemblies. To our knowledge, this is the first tool which is entirely based on this level of biological organization. SynTracker’s attractive features include that it does not require pre existing databases, and has a minimal sensitivity to sequencing errors and natural variation in microbial populations. SynTracker performs well when classifying isolate genomes and when tracking strains in longitudinal metagenomes. SynTracker could be used as a standalone tool or combined with existing tools in a multi-tool pipeline setup.

SynTracker is available at: https://github.com/leylabmpi/SynTracker

## Acknowledgements

We thank Nick Youngblut for providing comments on a previous version of this work.

## Methods

### SynTracker Pipeline

The SynTracker pipeline consists of three main parts. In the first part, SynTracker accepts a collection of reference genomes (a single genome per species), either fully assembled or as a collection of contigs. Each per-species reference is fragmented into a collection of 1kbp central-regions, which are binned and stored together.

In the second part SynTracker creates a blast Database, based on a user-provided collection of metagenomic assemblies or genomes. Next, it performs a blast search, for each of the central regioins, against the newly created blast database with a minimal identity of 97% and a minimal query coverage of 70%, i.e., 700bp. In the final step of this part, hits for each blast search are retrieved, using the *blastcmddb* command, in addition to a 2kbp region on each side of the blast hit. Hits with <2kbp both downstream and downstream to the hit are excluded from further analysis. Each retrieved sequence is denoted by its sample of origin and matching region in the reference genome.

In the third part of the pipeline genomic fragments are grouped by their matching region in the reference genome, and pairwise alignment is conducted to identify synteny blocks in each pair of sequences. The identification of synteny blocks in each pairwise alignment is performed using the “FindSynteny” function, in the “DECIPHER” R package (Wright 2016), with parameters “maxGap” and “maxSep” both set to 15. Additionally, only pairwise comparisons with a minimal overlap 4800 bp are considered for downstream analysis. Next, per each pairwise alignment, a synteny score is calculated, as described in equation 1:

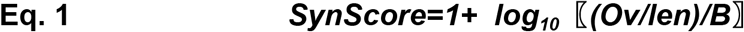

Where *Ov* stands for the accumulative length of the overlapping synteny blocks identified in the pairwise alignment, *len* denotes the length of the shorter sequence in each pair and *B* stands for the number of synteny blocks identified in each pairwise alignment.

In the final step of the third part of the pipeline, for each reference genome *n* genomic regions are randomly selected, per pair of metagenomic samples or genomes. APSS (average pairwise synteny scores) are calculated by averaging the individual pairwise synteny scores. Pairs of samples/genomes with <*n* regions are excluded from downstream analysis.

### In-silico evolutionary simulations

Calculation of the synteny scores per group of sampled cells was performed as described above, however, as the length of the genomic fragments used in the simulation was limited to 20kbp, synteny scores were based on a single alignment of the ~20kbp region, per pair of simulated genomes.

### Classification of bacterial genomes

Calculation of APSS values for *E.Coli* strain pairs was performed as described above and using the *E.coli* str. K-12 substr. MG1655 genome as a reference (NCBI Reference Sequence: NC_000913.3).

Phylogenetic trees were generated by conversion of APSS values to synteny distances, which equal to 1-APSS. All pairwise synteny distances were placed in a symmetric matrix which was used to calculate UPGMA phylogenetic trees, by employing the “phangorn” R-package (Schliep 2011).

### Tracking within-person strains

Longitudinal metagenomes were obtained from the NCBI-SRA database, and were quality filtered as described previously (Youngblut et al. 2020). Metagenomic samples were de-novo assembled using metaSPades (Nurk et al. 2017), with a maximal number of 20M reads/sample. ROC curves and matching APSS thresholds for each combination of species and sampling depth, in the testing set, were calculated using the R-programing language “pROC*”* package (Robin et al. 2011).

### Mother-infant strain transmission

Metagenomic samples were downloaded from the NCBI-SRA database, and were quality filtered and assembled as described above, however, as for some samples only one of the two matching read files passed our quality filtration, we performed the metagenomic assembly using single-end reads.

---

**Table S1:**
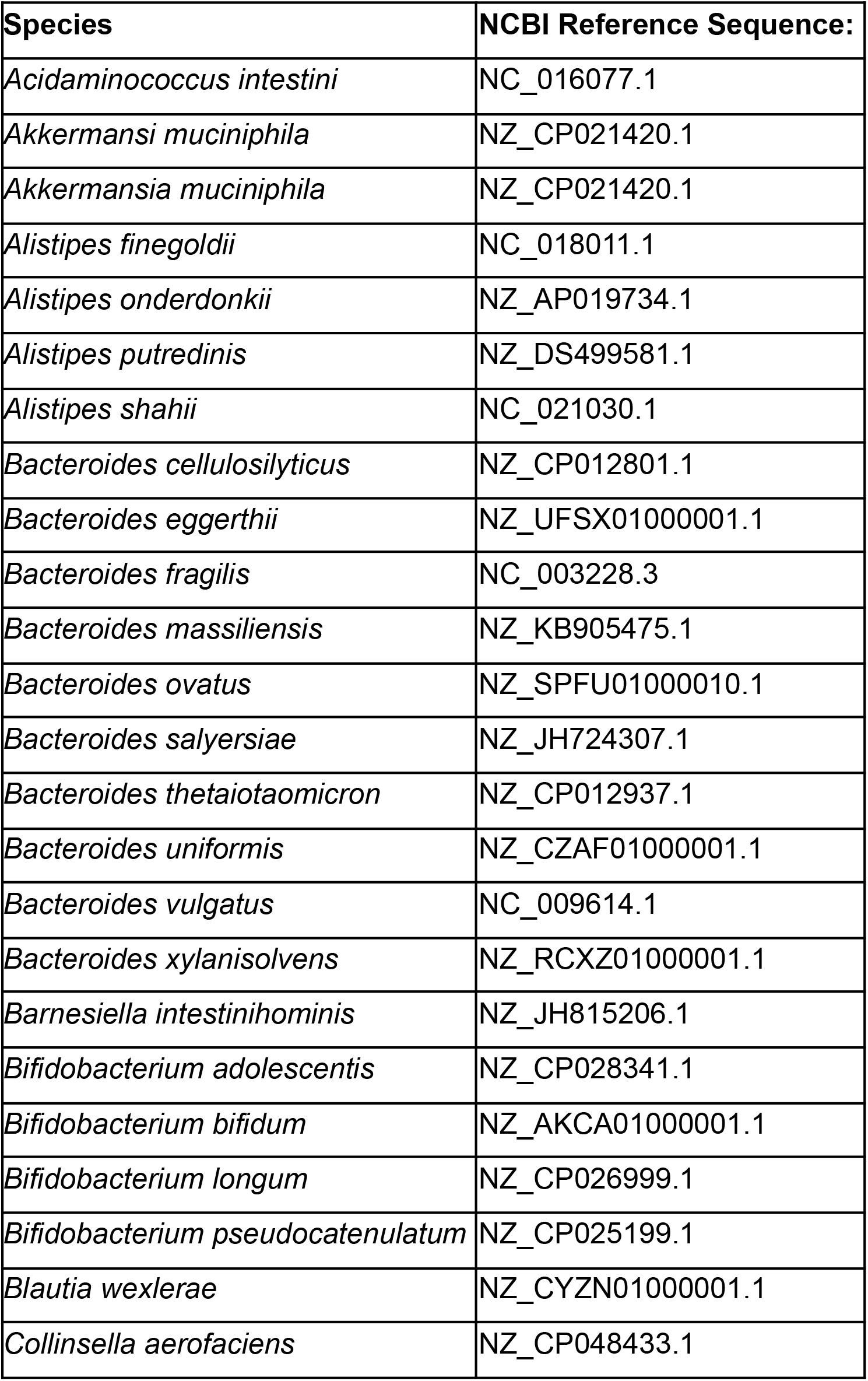
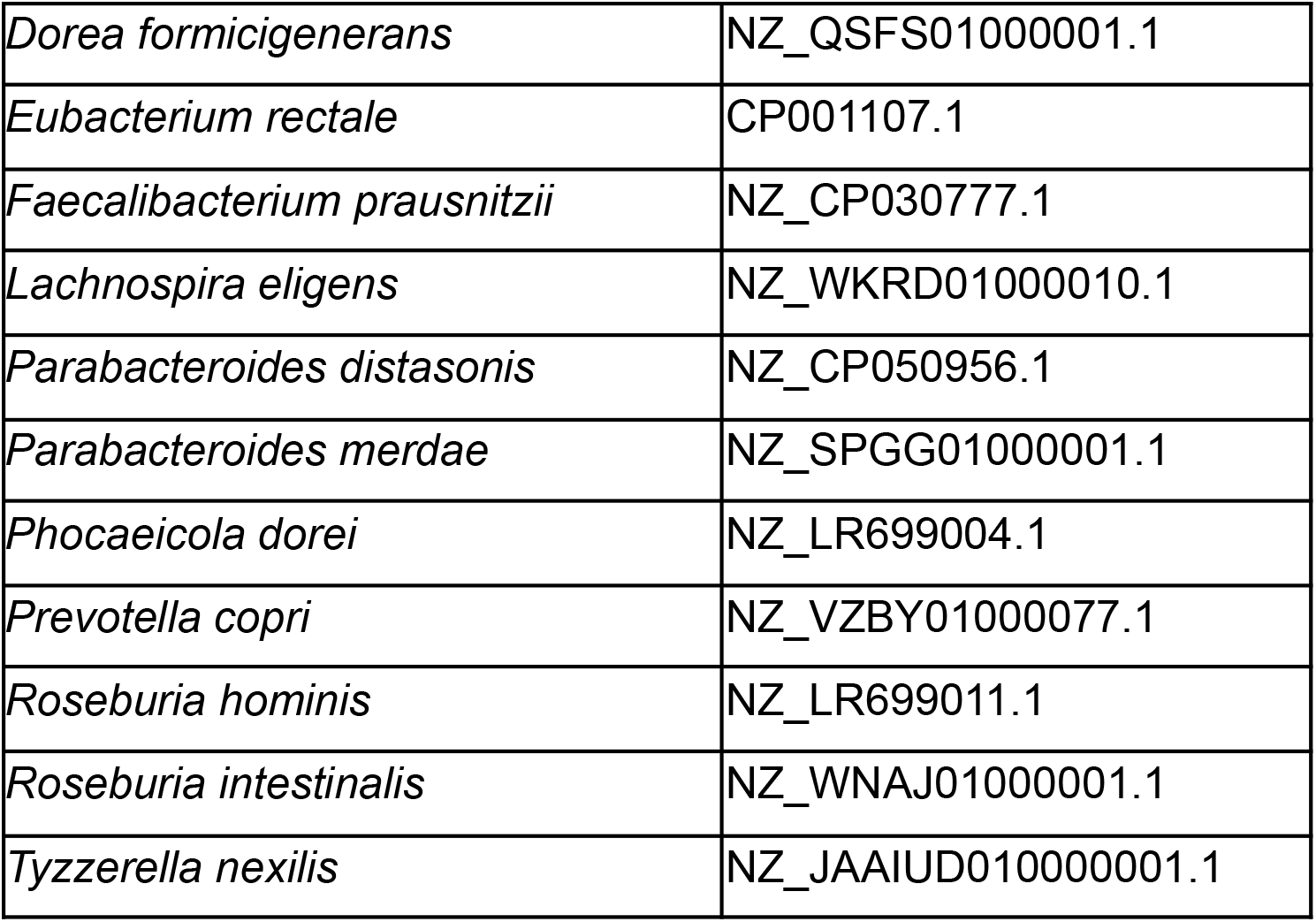
Reference genomes used in metagenomic analyses.

**Table S2:** Metagenomic samples used to evaluate the performance of SynTracker.

| ***NCBI Biosample*** | ***Individual*** | ***Sample name*** | ***set*** |
| --- | --- | --- | --- |
| SAMN11950002 | ab | ab-0140_MG | Testing |
| SAMN11950003 | ab | ab-0168_MG | Testing |
| SAMN11950006 | ad | ad-0002_MG | Testing |
| SAMN11950007 | ad | ad-0005_MG | Testing |
| SAMN11950014 | ae | ae-0008_MG | Testing |
| SAMN11950017 | ae | ae-0011_MG | Testing |
| SAMN11950025 | ae | ae-0022_MG | Testing |
| SAMN11950026 | ae | ae-0024_MG | Testing |
| SAMN11950029 | ae | ae-0027_MG | Testing |
| SAMN11950031 | ae | ae-0029_MG | Testing |
| SAMN11950047 | ae | ae-0052_MG | Testing |
| SAMN11950063 | ae | ae-0073_MG | Testing |
| SAMN11950069 | ag | ag-0001_MG | Testing |
| SAMN11950070 | ag | ag-0005_MG | Testing |
| SAMN11950073 | ai | ai-0002_MG | Testing |
| SAMN11950074 | ai | ai-0019_MG | Testing |
| SAMN11950077 | ak | ak-0001_MG | Testing |
| SAMN11950078 | ak | ak-0010_MG | Testing |
| SAMN11950107 | am | am-0027_MG | Testing |
| SAMN11950123 | am | am-0061_MG | Testing |
| SAMN11950159 | am | am-0097_MG | Testing |
| SAMN11950172 | am | am-0110_MG | Testing |
| SAMN11950229 | am | am-0172_MG | Testing |
| SAMN11950234 | am | am-0177_MG | Testing |
| SAMN11950254 | am | am-0199_MG | Testing |
| SAMN11950261 | am | am-0206_MG | Testing |
| SAMN11950297 | an | an-0017_MG | Testing |
| SAMN11950299 | an | an-0023_MG | Testing |
| SAMN11950322 | an | an-0051_MG | Testing |
| SAMN11950332 | an | an-0063_MG | Testing |
| SAMN11950339 | an | an-0071_MG | Testing |
| SAMN11950341 | an | an-0073_MG | Testing |
| SAMN11950354 | ao | ao-0007_MG | Testing |
| SAMN11950356 | ao | ao-0009_MG | Testing |
| SAMN11950366 | ao | ao-0026_MG | Testing |
| SAMN11950391 | ao | ao-0057_MG | Testing |
| SAMN11950402 | ao | ao-0069_MG | Testing |
| SAMN11950403 | ao | ao-0070_MG | Testing |
| SAMN11950406 | ao | ao-0073_MG | Testing |
| SAMN11950414 | ao | ao-0081_MG | Testing |
| SAMN11950466 | bp | bp-0002_MG | Testing |
| SAMN11950467 | bp | bp-0004_MG | Testing |
| SAMN11950472 | bs | bs-0001_MG | Testing |
| SAMN11950473 | bs | bs-0008_MG | Testing |
| SAMN11950488 | ca | ca-0001_MG | Testing |
| SAMN11950489 | ca | ca-0012_MG | Testing |
| SAMN11950500 | cg | cg-0001_MG | Testing |
| SAMN11950501 | cg | cg-0014_MG | Testing |
| SAMN11950506 | cj | cj-0001_MG | Testing |
| SAMN11950507 | cj | cj-0004_MG | Testing |
| SAMN11950523 | ct | ct-0001_MG | Testing |
| SAMN11950524 | ct | ct-0005_MG | Testing |
| SAMN11950527 | cv | cv-0001_MG | Testing |
| SAMN11950528 | cv | cv-0018_MG | Testing |
| SAMN11950551 | dg | dg-0001_MG | Testing |
| SAMN11950552 | dg | dg-0008_MG | Testing |
| SAMN11950555 | di | di-0001_MG | Testing |
| SAMN11950556 | di | di-0009_MG | Testing |
| SAMN11950559 | dk | dk-0001_MG | Testing |
| SAMN11950560 | dk | dk-0003_MG | Testing |
| SAMN11950513 | cn | cn-0006_MG | Testing |
| SAMN11950512 | cn | cn-0002_MG | Testing |
| SAMN11950511 | cm | cm-0022_MG | Testing |
| SAMN11950510 | cm | cm-0001_MG | Testing |
| SAMN11950509 | ck | ck-0028_MG | Testing |
| SAMN11950508 | ck | ck-0001_MG | Testing |
| SAMN11950499 | cf | cf-0037_MG | Testing |
| SAMN11950498 | cf | cf-0001_MG | Testing |
| SAMN11950497 | ce | ce-0007_MG | Testing |
| SAMN11950496 | ce | ce-0001_MG | Testing |
| SAMN11950495 | cd | cd-0050_MG | Testing |
| SAMN11950494 | cd | cd-0001_MG | Testing |
| SAMN11950485 | by | by-0059_MG | Testing |
| SAMN11950484 | by | by-0002_MG | Testing |
| SAMN11950483 | bx | bx-0057_MG | Testing |
| SAMN11950482 | bx | bx-0001_MG | Testing |
| SAMN11950463 | bn | bn-0038_MG | Testing |
| SAMN11950462 | bn | bn-0002_MG | Testing |
| SAMN11950461 | bm | bm-0013_MG | Testing |
| SAMN11950460 | bm | bm-0002_MG | Testing |
| SAMN11950459 | bl | bl-0009_MG | Testing |
| SAMN11950458 | bl | bl-0003_MG | Testing |
| SAMN11950457 | bk | bk-0025_MG | Testing |
| SAMN11950456 | bk | bk-0002_MG | Testing |
| SAMN11950455 | bi | bi-0026_MG | Testing |
| SAMN11950454 | bi | bi-0001_MG | Testing |
| SAMN11950453 | bh | bh-0112_MG | Testing |
| SAMN11950450 | be | be-0085_MG | Testing |
| SAMN11950449 | be | be-0001_MG | Testing |
| SAMN11950448 | bd | bd-0033_MG | Testing |
| SAMN11950447 | ba | ba-0002_MG | Testing |
| SAMN11950446 | ba | ba-0001_MG | Testing |
| SAMN11950443 | ay | ay-0058_MG | Testing |
| SAMN11950442 | ay | ay-0001_MG | Testing |
| SAMN11950441 | ax | ax-0090_MG | Testing |
| SAMN11950440 | ax | ax-0001_MG | Testing |
| SAMN11950439 | aw | aw-0036_MG | Testing |
| SAMN11950438 | aw | aw-0001_MG | Testing |
| SAMN11950433 | at | at-0044_MG | Testing |
| SAMN11950432 | at | at-0004_MG | Testing |
| SAMN11950431 | as | as-0033_MG | Testing |
| SAMN11950430 | as | as-0002_MG | Testing |
| SAMN11950427 | aq | aq-0015_MG | Testing |
| SAMN11950426 | aq | aq-0004_MG | Testing |
| SAMN11950425 | ap | ap-0037_MG | Testing |
| SAMN11950424 | ap | ap-0001_MG | Testing |
| SAMN11950000 | aa | aa-0154_MG | Training |
| SAMN11950001 | aa | aa-0163_MG | Training |
| SAMN11950004 | ac | ac-0002_MG | Training |
| SAMN11950005 | ac | ac-0038_MG | Training |
| SAMN11950008 | ae | ae-0001_MG | Training |
| SAMN11950012 | ae | ae-0005_MG | Training |
| SAMN11950019 | ae | ae-0014_MG | Training |
| SAMN11950023 | ae | ae-0020_MG | Training |
| SAMN11950027 | ae | ae-0025_MG | Training |
| SAMN11950028 | ae | ae-0026_MG | Training |
| SAMN11950032 | ae | ae-0030_MG | Training |
| SAMN11950067 | af | af-0003_MG | Training |
| SAMN11950068 | af | af-0060_MG | Training |
| SAMN11950071 | ah | ah-0002_MG | Training |
| SAMN11950072 | ah | ah-0028_MG | Training |
| SAMN11950075 | aj | aj-0001_MG | Training |
| SAMN11950076 | aj | aj-0061_MG | Training |
| SAMN11950079 | al | al-0002_MG | Training |
| SAMN11950080 | al | al-0025_MG | Training |
| SAMN11950135 | am | am-0073_MG | Training |
| SAMN11950158 | am | am-0096_MG | Training |
| SAMN11950180 | am | am-0118_MG | Training |
| SAMN11950190 | am | am-0128_MG | Training |
| SAMN11950238 | am | am-0181_MG | Training |
| SAMN11950250 | am | am-0195_MG | Training |
| SAMN11950262 | am | am-0207_MG | Training |
| SAMN11950288 | an | an-0002_MG | Training |
| SAMN11950295 | an | an-0015_MG | Training |
| SAMN11950301 | an | an-0026_MG | Training |
| SAMN11950303 | an | an-0028_MG | Training |
| SAMN11950334 | an | an-0065_MG | Training |
| SAMN11950336 | an | an-0067_MG | Training |
| SAMN11950346 | an | an-0080_MG | Training |
| SAMN11950347 | an | an-0081_MG | Training |
| SAMN11950358 | ao | ao-0011_MG | Training |
| SAMN11950359 | ao | ao-0012_MG | Training |
| SAMN11950392 | ao | ao-0058_MG | Training |
| SAMN11950393 | ao | ao-0059_MG | Training |
| SAMN11950404 | ao | ao-0071_MG | Training |
| SAMN11950405 | ao | ao-0072_MG | Training |
| SAMN11950418 | ao | ao-0085_MG | Training |
| SAMN11950470 | br | br-0001_MG | Training |
| SAMN11950471 | br | br-0002_MG | Training |
| SAMN11950480 | bw | bw-0001_MG | Training |
| SAMN11950481 | bw | bw-0033_MG | Training |
| SAMN11950492 | cc | cc-0002_MG | Training |
| SAMN11950493 | cc | cc-0023_MG | Training |
| SAMN11950502 | ch | ch-0001_MG | Training |
| SAMN11950503 | ch | ch-0008_MG | Training |
| SAMN11950514 | cp | cp-0001_MG | Training |
| SAMN11950515 | cp | cp-0009_MG | Training |
| SAMN11950525 | cu | cu-0001_MG | Training |
| SAMN11950526 | cu | cu-0009_MG | Training |
| SAMN11950541 | db | db-0001_MG | Training |
| SAMN11950542 | db | db-0015_MG | Training |
| SAMN11950553 | dh | dh-0001_MG | Training |
| SAMN11950554 | dh | dh-0010_MG | Training |
| SAMN11950557 | dj | dj-0001_MG | Training |
| SAMN11950558 | dj | dj-0016_MG | Training |
| SAMN11950561 | dl | dl-0001_MG | Training |
| SAMN11950562 | dl | dl-0006_MG | Training |
| SAMN11950517 | cq | cq-0035_MG | Training |
| SAMN11950516 | cq | cq-0001_MG | Training |
| SAMN11950505 | ci | ci-0052_MG | Training |
| SAMN11950504 | ci | ci-0001_MG | Training |
| SAMN11950491 | cb | cb-0051_MG | Training |
| SAMN11950490 | cb | cb-0001_MG | Training |
| SAMN11950487 | bz | bz-0033_MG | Training |
| SAMN11950486 | bz | bz-0001_MG | Training |
| SAMN11950479 | bv | bv-0024_MG | Training |
| SAMN11950478 | bv | bv-0001_MG | Training |
| SAMN11950477 | bu | bu-0080_MG | Training |
| SAMN11950476 | bu | bu-0001_MG | Training |
| SAMN11950475 | bt | bt-0039_MG | Training |
| SAMN11950474 | bt | bt-0001_MG | Training |
| SAMN11950469 | bq | bq-0068_MG | Training |
| SAMN11950468 | bq | bq-0002_MG | Training |
| SAMN11950465 | bo | bo-0122_MG | Training |
| SAMN11950464 | bo | bo-0001_MG | Training |
| SAMN11950452 | bf | bf-0108_MG | Training |
| SAMN11950451 | bf | bf-0003_MG | Training |
| SAMN11950445 | az | az-0036_MG | Training |
| SAMN11950444 | az | az-0001_MG | Training |
| SAMN11950437 | av | av-0107_MG | Training |
| SAMN11950436 | av | av-0006_MG | Training |
| SAMN11950435 | au | au-0066_MG | Training |
| SAMN11950434 | au | au-0002_MG | Training |
| SAMN11950429 | ar | ar-0039_MG | Training |
| SAMN11950428 | ar | ar-0002_MG | Training |
| SAMN11950423 | ao | ao-0090_MG | Training |
| SAMN11950350 | ao | ao-0001_MG | Training |
| SAMN11950349 | an | an-0083_MG | Training |
| SAMN11950287 | an | an-0001_MG | Training |
| SAMN11950286 | am | am-0231_MG | Training |
| SAMN11950116 | am | am-0054_MG | Training |
| SAMN11950550 | df | df-0030_MG | Training |
| SAMN11950549 | df | df-0001_MG | Training |
| SAMN11950548 | de | de-0031_MG | Training |
| SAMN11950547 | de | de-0001_MG | Training |
| SAMN11950546 | dd | dd-0041_MG | Training |
| SAMN11950545 | dd | dd-0001_MG | Training |
| SAMN11950544 | dc | dc-0028_MG | Training |
| SAMN11950543 | dc | dc-0001_MG | Training |
| SAMN11950540 | da | da-0044_MG | Training |
| SAMN11950539 | da | da-0001_MG | Training |
| SAMN11950538 | cz | cz-0039_MG | Training |
| SAMN11950537 | cz | cz-0001_MG | Training |
| SAMN11950536 | cy | cy-0037_MG | Training |
| SAMN11950534 | cy | cy-0001_MG | Training |
| SAMN11950533 | cx | cx-0014_MG | Training |
| SAMN11950532 | cx | cx-0001_MG | Training |
| SAMN11950531 | cw | cw-0053_MG | Training |
| SAMN11950529 | cw | cw-0001_MG | Training |
| SAMN11950522 | cs | cs-0029_MG | Training |
| SAMN11950520 | cs | cs-0001_MG | Training |
| SAMN11950519 | cr | cr-0043_MG | Training |
| SAMN11950518 | cr | cr-0001_MG | Training |

